# Non-cell autonomous cardiomyocyte regulation complicates gene supplementation therapy for *LMNA* cardiomyopathy

**DOI:** 10.1101/2023.07.18.549413

**Authors:** Yueshen Sun, Congting Guo, Zhan Chen, Junsen Lin, Luzi Yang, Yueyang Zhang, Chenyang Wu, Dongyu Zhao, Blake Jardin, William T. Pu, Mingming Zhao, Erdan Dong, Xiaomin Hu, Shuyang Zhang, Yuxuan Guo

**Affiliations:** Department of Cardiology, Peking Union Medical College Hospital, Chinese Academy of Medical Science & Peking Union Medical College, 100730, Beijing, China; School of Basic Medical Sciences, Peking University Health Science Center, Beijing, 100191, China; Peking University Institute of Cardiovascular Sciences, Beijing, 100191, China; Department of Biomedical informatics, School of Basic Medical Sciences, Peking University, Beijing, 100191, China; State Key Laboratory of Vascular Homeostasis and Remodeling, Peking University, Beijing, 100191, China; Department of Cardiology, Boston Children’s Hospital, Boston, MA 02115, USA; Harvard Stem Cell Institute, Cambridge, MA 02138, USA; Department of Cardiology and Institute of Vascular Medicine, Peking University Third Hospital, Beijing, 100191, China; Research Unit of Medical Science Research Management/Basic and Clinical Research of Metabolic Cardiovascular Diseases, Chinese Academy of Medical Sciences, Haihe Laboratory of Cell Ecosystem, Beijing, 100191, China; National Health Commission Key Laboratory of Cardiovascular Molecular Biology and Regulatory Peptides, Beijing, 100191, China; Beijing Key Laboratory of Cardiovascular Receptors Research, Beijing, 100191, China; Research Center for Cardiopulmonary Rehabilitation, University of Health and Rehabilitation Sciences Qingdao Hospital (Qingdao Municipal Hospital), School of Health and Life Sciences, University of Health and Rehabilitation Sciences; Qingdao 266071, China; Department of Medical Research Center, Peking Union Medical College Hospital, Chinese Academy of Medical Science and Peking Union Medical College, Beijing, China, 100730, Beijing, China; State Key Laboratory of Complex Severe and Rare Diseases, Peking Union Medical College Hospital, Chinese Academy of Medical Science and Peking Union Medical College, 100730, Beijing, China

**Keywords:** *LMNA* cardiomyopathy, gene therapy, adeno-associated virus, non-cell autonomous, cardiomyocyte maturation

## Abstract

**Aims:** Recombinant adeno-associated viruses (rAAVs) are federally approved gene delivery vectors for in vivo gene supplementation therapy. Loss-of-function truncating variants of *LMNA*, the coding gene for Lamin-A/C, are one of the primary causes of inherited dilate cardiomyopathy (DCM). Here we aim to study whether AAV-based *LMNA* supplementation could treat *LMNA* deficiency-triggered cardiac defects.

**Methods and Results:** We compared whole-body, cardiomyocyte-specific and genetic-mosaic mouse models that carry *Lmna* truncating variants at the same genetic loci and uncovered primarily a non-cell autonomous impact of *Lmna* on cardiomyocyte maturation. Whole-body lamin-A supplementation by rAAVs moderately rescued the cardiac defects in *Lmna* germline mutants. By contrast, cardiomyocyte-specific lamin-A addback failed to restore the cardiomyocyte growth defects. A Cre-loxP-based AAV vector that expresses lamin-A throughout the body but excluding the heart was able to restore cardiomyocyte growth in *Lmna* germline mutants.

**Conclusions:** *Lmna* regulates cardiomyocyte growth non-cell autonomously. Non-myocytes are the key cell targets for a successful gene therapy for *LMNA*-associated cardiac defects.

**Translational perspective:** *LMNA* truncating mutations are among the major causes of inherited DCM. AAV gene supplementation therapy is emerging as a promising strategy to treat genetic cardiomyopathy, but whether this strategy is suitable for *LMNA* cardiomyopathy remained unclear. Our study counterintuitively showed that the cardiomyocytes are not necessarily the correct therapeutic cell targets for AAV-based treatment of *LMNA* cardiomyopathy. By contrast, careful elucidation of cell-autonomous versus non-cell-autonomous gene functions is essential for the proper design of a gene supplementation therapy for cardiomyopathy.

**Graphical Abstract:** 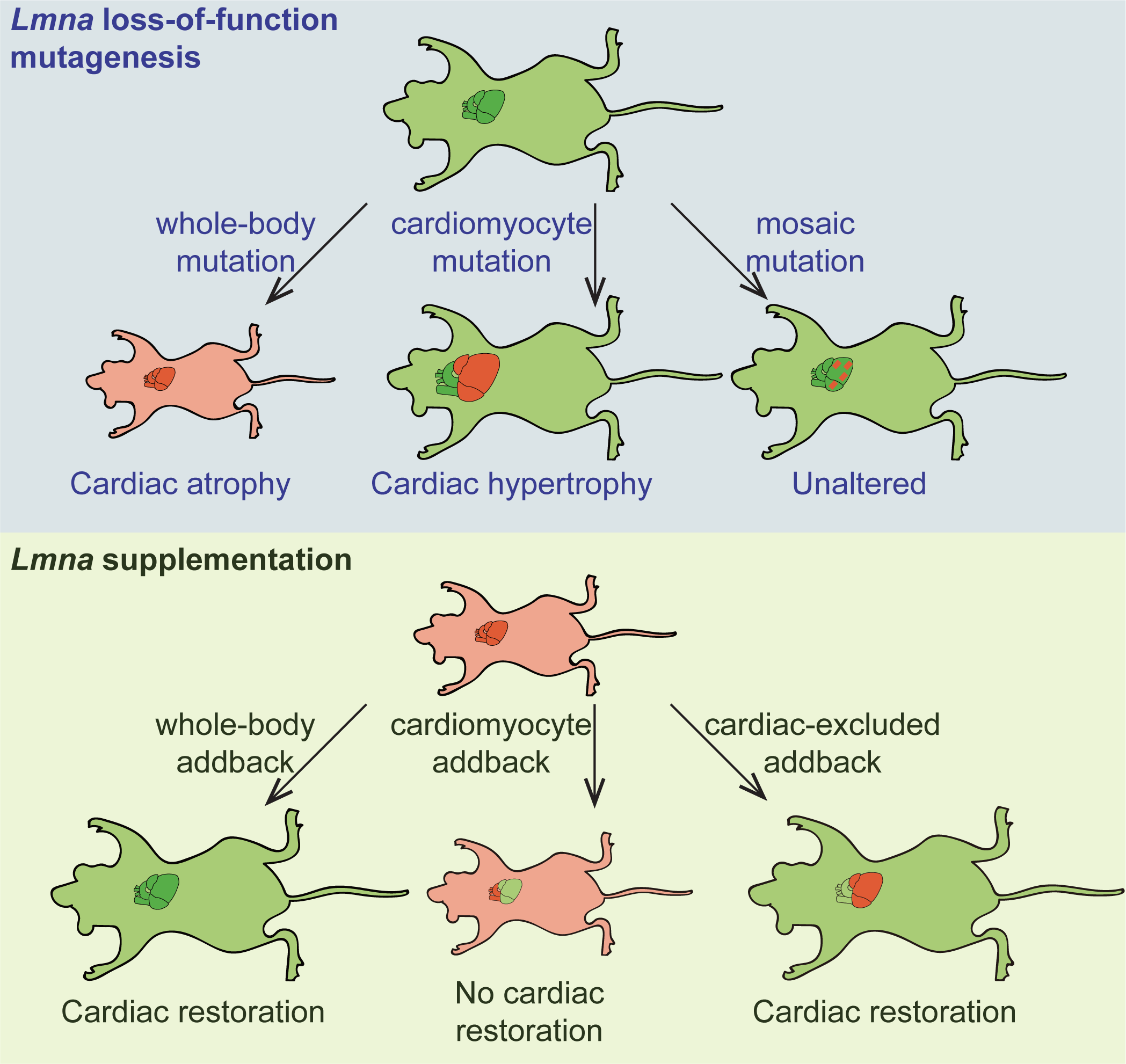

## Introduction

Recombinant adeno-associated viruses (rAAVs) are popular gene delivery vectors for in vivo gene therapy^1^. The federally approved applications of rAAVs mainly involve gene supplementation therapy, in which rAAVs deliver transgenes to replace or supplement endogenous genes that harbor disease-causing loss-of-function (LOF) mutations^2^. A successful AAV therapy requires rAAVs to efficiently deliver the therapeutic transgene to the correct cell and tissue targets. Myocardium is among the primary tissues that can be efficiently transduced by rAAVs^3^. In animal models, rAAV-based gene supplementation therapy has successfully alleviated or reversed cardiac malfunctioning in an array of genetic diseases that are associated with myocardium defects, such as the Danon disease^4^, Barth syndrome^5^ and arrythmogenic cardiomyopathy^6^.

Dilated cardiomyopathy (DCM) is one of the most prevalent forms of cardiomyopathy that deposit a high risk for heart failure and sudden cardiac death^7^. Up to 30%-50% DCM cases are associated with genetic factors, particularly including the truncating variants of the top-two DCM-causing genes TTN and *LMNA*^8^. TTN is an enormous gene that is too large to be delivered by rAAVs, but the coding sequences in *LMNA* are less than 2 kilobases (kb), which ideally fits into the 4.7kb size limit of rAAV genome. Therefore, *LMNA* appears to be an attractive gene target in rAAV gene therapy for DCMs that are caused by *LMNA* deficiency.

The gene supplementation therapy for *LMNA*-associated cardiomyopathy is desirable also because of the complicated pathogenic mechanisms downstream of *LMNA* mutations. *LMNA* encodes A-type lamins, primarily lamin-A and lamin-C (collectively called Lamin-A/C), which are type-V intermediate filament proteins constituting the nuclear lamina^9, 10^. LOF mutations of *LMNA* are known to trigger many diverse defects in cytoskeleton organization^11^, nuclear mechanics^12^, signal transduction^13^, DNA damage responses^14, 15^, chromatin organization^16^, epigenetic landscape^17^ and transcriptional regulation^13^. Therefore, it is very difficult to identify one universal therapeutic target to treat all defects that are associated with *LMNA*-associated cardiomyopathy. Instead, AAV-based supplementation of *LMNA* gene appears to be a more realistic approach to mitigate the disease phenotypes, although no such gene therapy studies have been reported.

Unlike many cardiomyopathy-causing genes that are restrictively expressed in the heart, *LMNA* is widely expressed in many differentiated cell types^18^. Cardiomyocyte-specific ablation of *Lmna* in mice resulted in severe DCM phenotypes^19, 20^, supporting the classic notion that cardiomyocyte is the primary cell type that contributes to *LMNA* cardiomyopathy. However, in DCM patients, *LMNA* mutations are also expressed in non-myocytes throughout the body. Recent evidence starts to uncover the pathogenic role of non-myocytes in *LMNA* DCM pathogenesis^21, 22^. Thus, more extensive comparison of cardiomyocyte versus non-myocyte contributions to *LMNA* cardiomyopathy would be necessary to determine if cardiomyocytes are indeed the correct targets for gene supplementation therapy.

DCM is characterized by ventricular enlargement, eccentric cardiac hypertrophy and axial cardiomyocyte elongation, thus a key assessment parameter in the gene therapy for *LMNA*-associated DCM is relevant to cardiomyocyte size and shape. Unfortunately, the existing *LMNA* cardiomyopathy models have exhibited inconsistent or even seemingly contradictory phenotypes in cardiomyocyte hypertrophy. For example, in germline hypomorphic *Lmna* mutant mice, which are often denoted as the *Lmna*^−/−^ mice^23, 24^, cardiomyocytes were reported to exhibit an atrophic phenotype within two weeks after birth, which is characteristic of cardiomyocyte maturation defect^25, 26^. By contrast, in mice carrying cardiomyocyte-specific *Lmna* LOF mutations, although DCM phenotypes also arise during the key postnatal cardiac maturation phase, these mice developed elevated ventricular mass and cardiomyocyte hypertrophy^19, 20^. Interestingly, in studies using human induced pluripotent stem-cell derived cardiomyocytes (hiPSC-CMs) to model *LMNA* cardiomyopathy, neither atrophy nor hypertrophy phenotypes were reported yet, further questioning a direct role of *LMNA* in regulating cardiomyocyte morphology and cardiac remodeling.

We previously demonstrated that cardiomyocyte maturation and cardiac hypertrophy analysis can be confounded by non-cell autonomous regulation of cardiomyocytes^26, 27^. This problem can be solved by rAAV-based genetic mosaics analysis. This approach harnessed low-dose rAAV transduction into a small fraction of cardiomyocytes to circumvent secondary effects from whole-body or cardiac dysfunction ^26–30^. In this study, we generated cardiac genetic mosaic mouse models carrying *Lmna* truncating mutations to determine the cell autonomous versus non-cell autonomous *Lmna* functions in cardiomyocytes. Based on this analysis, we further investigated cardiomyocytes versus non-myocytes as the essential cell targets for an effective AAV *LMNA* supplementation therapy.

## Results

### A germline *Lmna* truncation perturbs cardiomyocyte growth while maintaining a normal systolic function

We introduced a 5-base-pair (5bp) deletion into *Lmna* exon10 (Fig.1A) via Cas9-mediated zygotic gene editing^31^ and denoted this mutation as *Lmna*^Δ^. *Lmna*^Δ^ localized to the last exon that encodes both lamin-A and lamin-C proteins, thus *Lmna*^Δ^ was expected to shift *Lmna* open reading frames, producing a protein truncated at C-terminus (denoted as Lamin-Δ) (Fig.1A). Real-time quantitative PCR (RT-qPCR) and RNA sequencing (RNA-Seq) analysis revealed reduced *Lmna* mRNA level in postnatal day 14 (P14) *Lmna* ^Δ/Δ^ hearts (Fig.1B), suggesting the activation of nonsense-mediated mRNA decay by *Lmna* truncating mutations as previously reported^32^. Western blot validated the depletion of Lamin-A/C proteins, which was replaced by the expression of the lamin-Δ protein (Fig.1C). Immunofluorescence analysis of lamin-A/C in cardiac cryo-sections also revealed dramatically reduced signals at the nuclear periphery in *Lmna*^Δ/Δ^ hearts (Fig.1D). The reduction and truncation of lamin-A/C proteins was also validated in *Lmna*^Δ/Δ^ liver (Fig. S1A). Thus, we successfully generated a new mouse model that globally harbors a *Lmna* truncating mutation.

**Figure 1.**
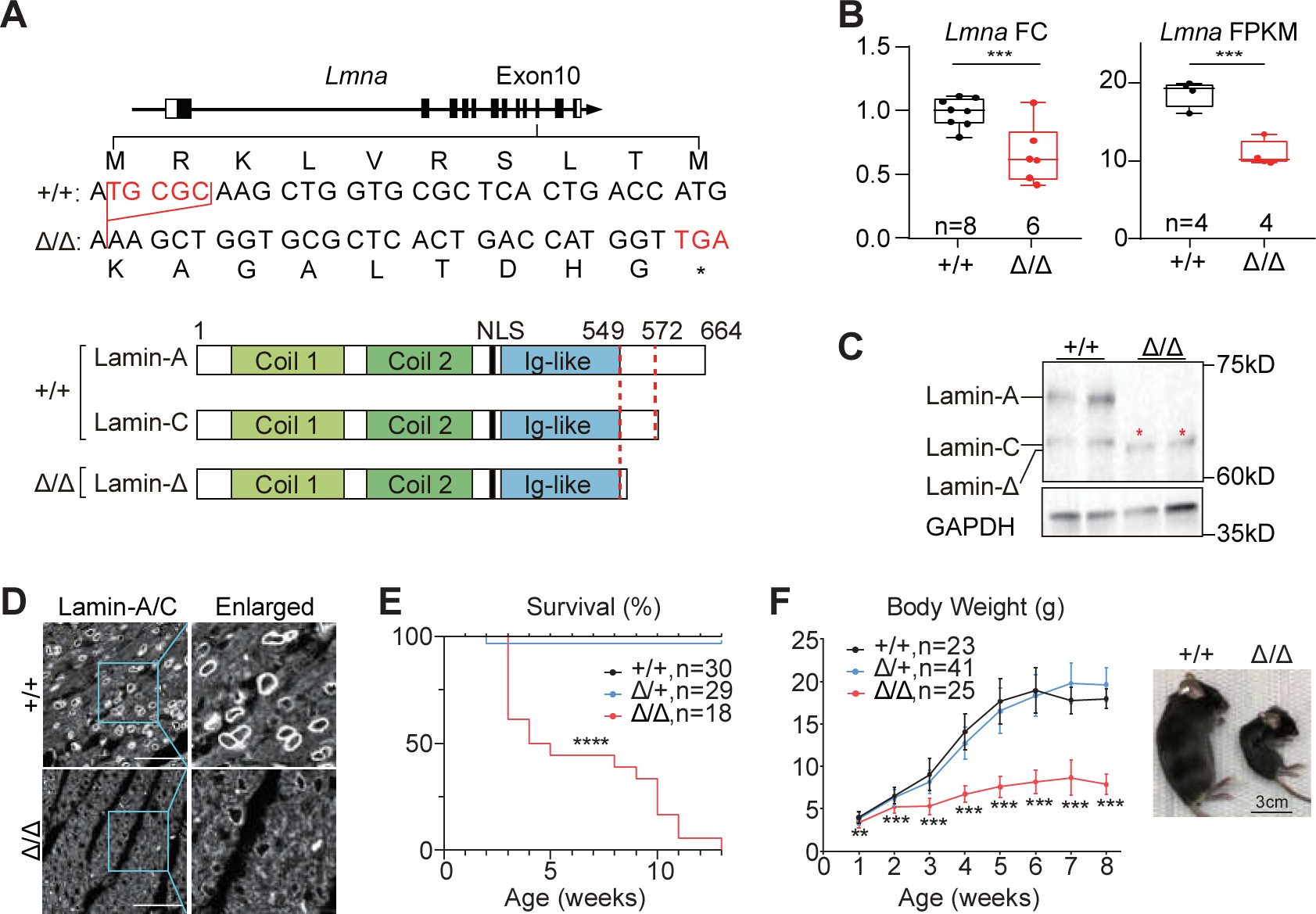
A new mouse model carrying a *Lmna* truncating mutation. **(A)** The top diagram showing a 5bp deletion in *Lmna* exon 10 in the *Lmna*^Δ^ allele and the introduction of a premature stop codon. The bottom diagram showing the resultant truncated protein by the mutation. **(B)** RT-qPCR and RNA-seq quantification of *Lmna* mRNA in P14 heart apexes. In box plots, horizontal lines indicate the median, 25th and 75th quantiles; whiskers extend to the extreme values; dots represent raw data points. **(C)** Western blot analysis of *Lmna*-expressed proteins in P14 heart apexes. Red star indicates the truncated protein bands. **(D)** Immunofluorescence of *Lmna*-expressed proteins in heart sections. Bars, 50μm. **(E-F)** Survival curve **(E)** and growth curve **(F)** of the mice. Mean±SD. Bright-field images of 6-week-old mice is presented to the right. Log-rank test was performed to analyze survival rates **(E)**. Student’s t-test in other panels. **p<0.01; ***p<0.001; ****p<0.0001.

*Lmna*^Δ/+^ intercross reproduced *Lmna*^+/+^, *Lmna*^Δ/+^, and *Lmna*^Δ/Δ^ pups following the Mendelian ratio at birth (Fig. S1B), but before the 13^th^ week after birth, all *Lmna*^Δ/Δ^ animals died (Fig. 1E). The body growth of *Lmna*^+/+^ and *Lmna*^Δ/+^ mice appeared indistinguishable, while *Lmna*^Δ/Δ^ pups exhibited severe retardation in gaining body weight (Fig. 1F), which agreed to the observation in previously reported *Lmna* hypomorphic mice^23, 24^. The heart of the *Lmna*^Δ/Δ^ pups also appeared smaller than their littermate controls (Fig. 2A). Consistent with this observation, echocardiogram analysis demonstrated reduced left ventricular diameters (LVIDs and LVIDd) and posterior wall thickness (LVPWs and LVPWd) in *Lmna*^Δ/Δ^ hearts (Fig. 2B). Strikingly, echocardiogram showed normal cardiac fractional shortening (FS) in *Lmna*^Δ/Δ^ mice at 2 and 6 weeks of age (Fig.2C). Picrosirius red staining detected no obvious changes in cardiac fibrosis (Fig. 2D) and RT-qPCR analysis showed no expression changes of the cardiac fibrosis marker *Ctgf* (Fig. 2E).

**Figure 2.**
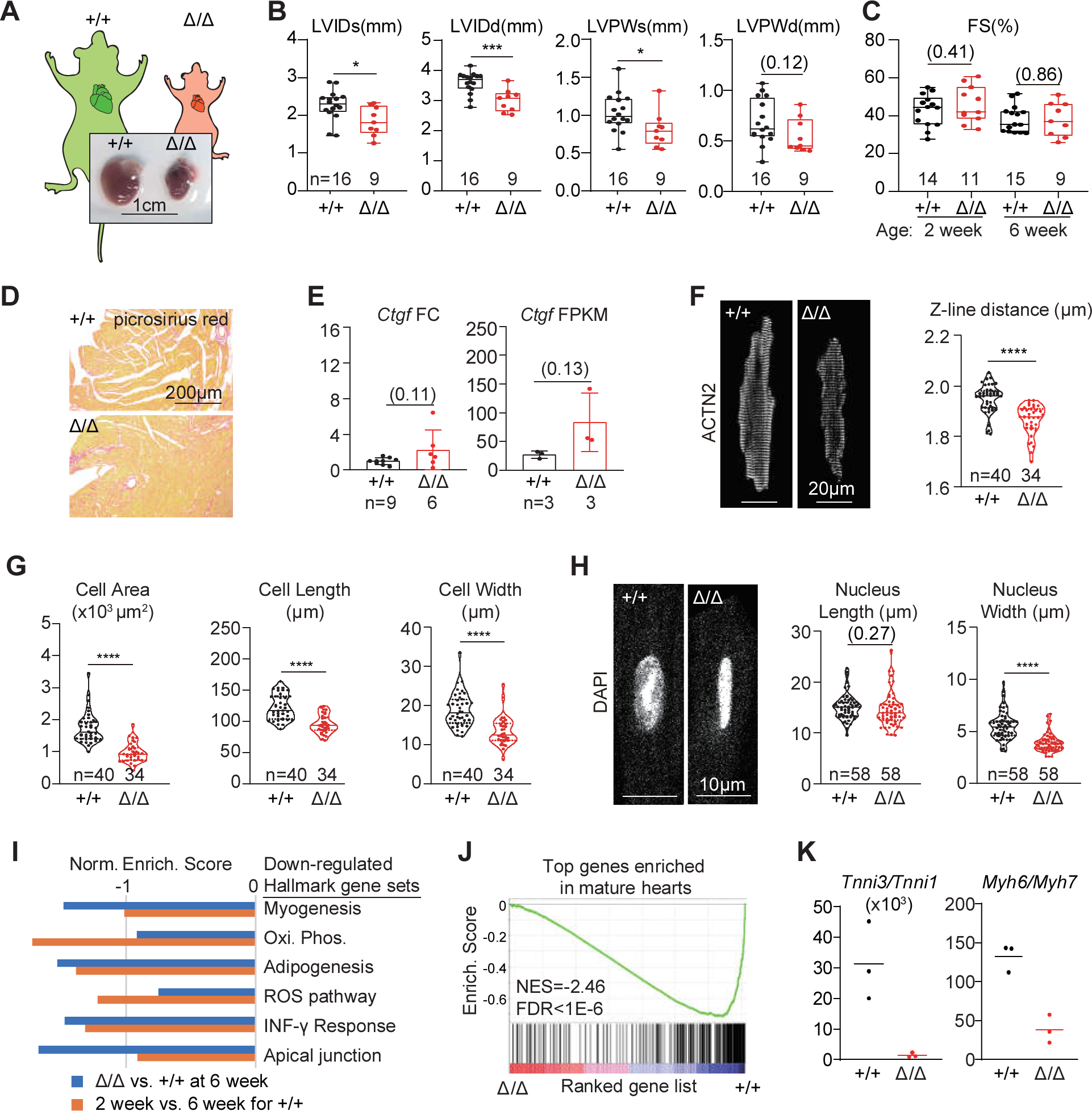
Cardiac maturation defects in *Lmna*^Δ/Δ^ mice. **(A)** Bright-field images of hearts extracted from 6-week-old mice. **(B-C)** Echocardiogram analysis of the 6-week-old mice. FS, fractional shortening; LVIDd, left ventricular internal diameter end diastole; LVIDs, left ventricular internal diameter end systole; LVPWd, left ventricular posterior wall end diastole; LVPWs, left ventricular systolic posterior wall end systole. **(D)** Picrosirius red staining on paraffin sections of the ventricles. **(E)** RT-qPCR and RNA-Seq analysis of Ctgf. **(F)** Immunostaining of ACTN2 on isolated cardiomyocytes and quantification of the distances between adjacent Z-lines. **(G)** Quantification of cardiomyocyte projected cell area, cell length and cell width. **(H)** DAPI-stained cell nuclei and quantification of nuclear length and width. In **(B-H)**, two-tailed t-test: *p<0.05; **p<0.01; ***p<0.001; non-significant P values in parentheses. **(I)** GSEA of the differentially expressed genes in RNA-Seq data. **(J)** Enrichment of cardiac mature marker among down-regulated genes in *Lmna* ^Δ/Δ^ ventricles. **(K)** Quantification of cardiomyocyte maturation isoform switching markers in RNA-Seq data.

Next we characterized the *Lmna*^Δ/Δ^ cardiac phenotypes at the cellular level by analyzing cardiomyocytes that were isolated via Langendorff collagenase perfusion. Immunostaining of α-actinin-2 (ACTN2), a marker for sarcomere Z-lines, revealed reduced sarcomere length in the mutant cardiomyocytes (Fig. 2F). These mutant cardiomyocytes appeared smaller in projected cell area, cell length and cell width (Fig. 2G). Agreeing to previous studies^33^, the size of *Lmna* ^Δ/Δ^ cardiomyocyte nuclei appeared smaller mainly because of the reduced nuclear width (Fig. 2H).

To further investigate the *Lmna*^Δ/Δ^ cardiac phenotypes at the gene expression level, we next performed RNA-seq differential expression analysis of 6-week *Lmna*^+/+^ versus *Lmna*^Δ/Δ^ heart ventricles (Fig. S2A-B, Table S1). As a reference data for postnatal heart maturation, we also conducted RNA-seq analysis and compared 2-week versus 6-week wildtype heart ventricles (Fig. S2C-D, Table S2). Gene set enrichment analysis (GSEA) using the Hallmark gene set^34^ uncovered significantly reduced myogenesis, oxidative phosphorylation and adipogenesis gene expression in the *Lmna*^Δ/Δ^ group (Fig. 2I). The same gene sets were also lower in the 2-week wildtype group (Fig. 2I), confirming that these gene sets were relevant to cardiac maturation^25, 35^. We next directly used the 2-week versus 6-week analysis to build a reference gene set of cardiac maturation markers, and performed GSEA on this gene set. This analysis clearly showed that, at the whole transcriptome level, mature heart markers were significantly reduced in *Lmna*^Δ/Δ^ hearts as compared to control hearts (Fig. 2J). Tnni3:Tnni1 and Myh6:Myh7 ratios, which are the classic cardiomyocyte maturation metrics^25, 35^, were also reduced in *Lmna*^Δ/Δ^ hearts (Fig. 2K). Together, these data showed that *Lmna*^Δ/Δ^ mice exhibited global cardiomyocyte maturation defects.

### Cardiomyocyte-specific *Lmna* truncation results in dilated cardiomyopathy and pathological hypertrophy

We reasoned that the *Lmna*^Δ/Δ^ mice exhibited the relatively normal heart function either because the *Lmna*^Δ^ mutation was not cardiac pathogenic, or the cardiac dysfunction was mitigated by the growth retardation of the whole body, which resulted in less demand for cardiac output. To distinguish between these two possibilities, we exploited a CRISPR/Cas9/AAV9-based somatic mutagenesis (CASAAV) system^27, 36^ to introduce *Lmna* ^Δ^-like mutations specifically in cardiomyocytes. In detail, we constructed an AAV9 vector that used U6 promoters to express sgRNAs targeting *Lmna* exon 10 at the same loci where the *Lmna*^Δ^ mutation was located (Fig. 3A-B). This vector also harbored a *Tnnt2* promoter (also called cTnT promoter)^37^ to express Cre recombinase specifically in cardiomyocytes. Therefore, when this AAV-*Tnnt2*-Cre-U6-*Lmna*-sgRNA (AAV-*Lmna*-sgRNA) vector was delivered into a Rosa^CAG-LSL-Cas9-tdTomato^ (Rosa^Cas9-Tom^) mice^36^, CRISPR/Cas9-based *Lmna* mutagenesis would selectively occur in cardiomyocytes that were labelled by tdTomato (Fig. 3A-C).

**Figure 3.**
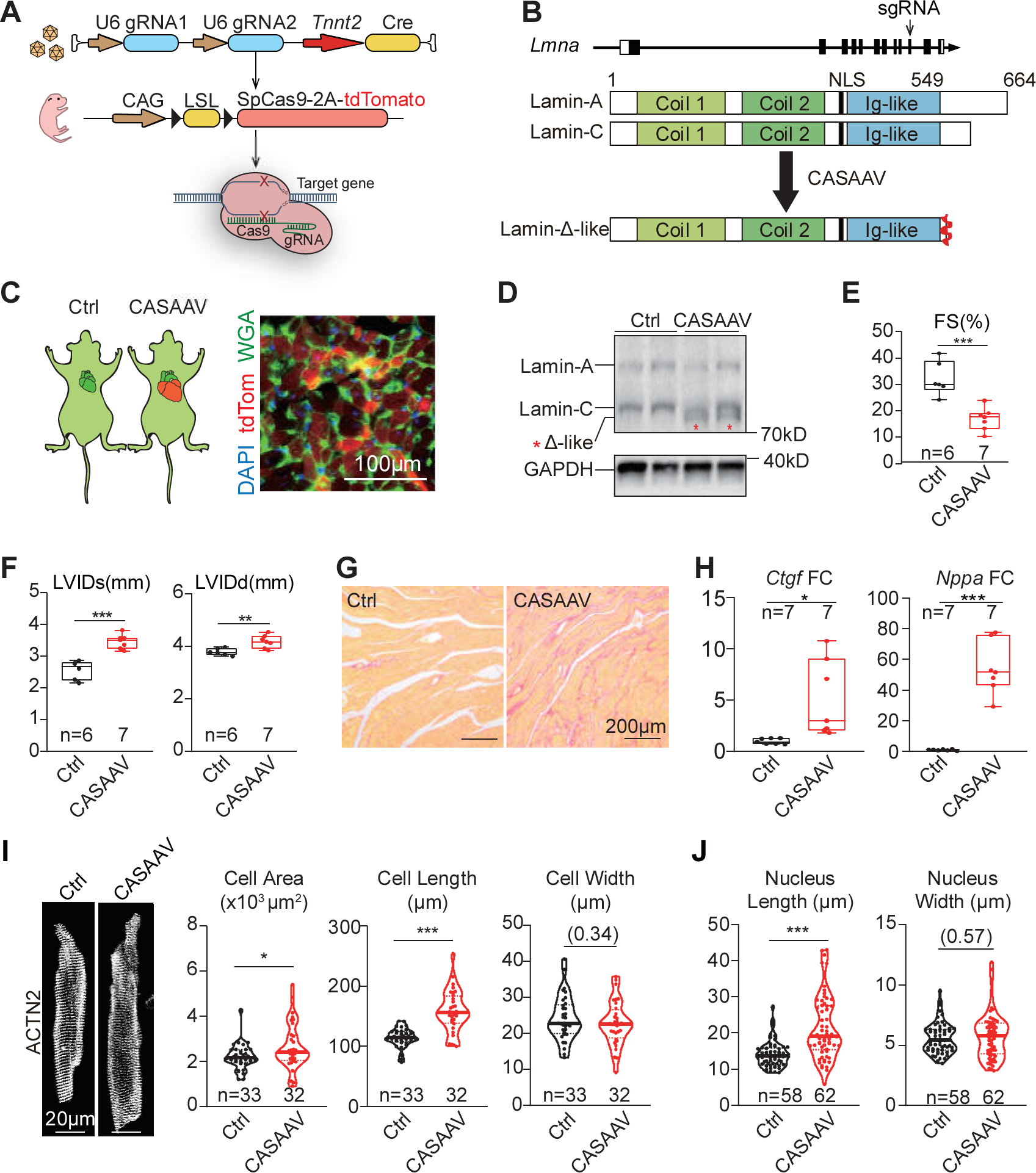
Cardiac hypertrophy in mice undergoing CASAAV-based cardiac mutagenesis on *Lmna*. **(A)** A diagram of CASAAV. **(B)** The expected gene expression consequence of CASAAV. **(C)** A representative fluorescence image of heart sections undergoing CASAAV. TdTomato labels AAV-transduced mutant cardiomyocytes. **(D)** Western blot analysis of *Lmna*-expressed proteins in P14 heart apexes. **(E-F)** Echocardiogram analysis of the 6-week-old mice. **(G)** Picrosirius red staining on 6-week-old heart sections. **(H)** RT-qPCR analysis of *Ctgf* and *Nppa*. **(I)** Immunostaining of ACTN2 on isolated cardiomyocytes and quantification of cardiomyocyte morphology. **(J)** Quantification of nuclear morphology. Two-tailed t-test: *p<0.05; ***p<0.001; non-significant P values in parentheses.

We first applied 2×10^11^ AAV-*Lmna*-sgRNA particles into P1 Rosa^Cas9-Tom^ mice and analyzed heart samples at 6 weeks after birth. Fluorescence imaging of cardiac cryo-sections detected tdTomato signals in up to 90% cardiomyocytes (Fig. 3C). As compared to AAV-*Tnnt2*-Cre treated control hearts, western blot detected the expression of truncated lamin-Δ-like proteins in the CASAAV-treated hearts at the molecular weight similar to the lamin-Δ protein in the *Lmna*^Δ/Δ^ hearts (Fig. 3D). Targeted high-throughput sequencing of the CASAAV-treated genomic regions confirmed the introduction of frame-shifting mutations similar to *Lmna*^Δ^ (Fig. S3).

In contrast to the relatively normal systolic function in *Lmna* ^Δ/Δ^ mice, CASAAV-mediated *Lmna* mutagenesis resulted in significant reduction of FS in the heart (Fig. 3E). LVIDd and LVIDs increased after CASAAV treatment for 6 weeks (Fig. 3F). CASAAV treatment also elevated picrosirius red-labeled cardiac interstitial fibrosis (Fig. 3G) and upregulated the expression of *Ctgf* and the cardiac stress marker *Nppa* in the heart (Fig. 3H). CASAAV treatment also increased projected cardiomyocyte area and cardiomyocyte length, while cell width was not affected (Fig. 3I), which was a characteristic cardiomyocyte elongation phenotype in DCM. Similarly, CASAAV-treated cell nuclei also become elongated (Fig. 3J). Together, this phenotypic comparison between whole-body and cardiomyocyte-specific *Lmna* mutagenesis demonstrates that the *Lmna* truncating mutation causes DCM only when the weakened heart function is insufficient to support a normally growing body.

### Cardiac genetic mosaic analysis uncovered non-cell autonomous regulation of cardiomyocyte morphology by *Lmna*

The seemingly contradictory cardiomyocyte morphology phenotypes in whole-body versus cardiomyocyte-specific *Lmna* mutants (Fig. 2G vs. Fig. 3I) suggested that the impact of *Lmna* mutation on cardiomyocyte morphology is secondary to the pathophysiological context. We next performed a cardiac genetic mosaic analysis^27, 28^ to test this hypothesis. We conducted a serial dilution of the AAV-*Lmna*-sgRNA vector and injected high (2×10^11^), mid (2×10^10^) and low (2×10^9^) doses of AAV into P1 Rosa^Cas9-Tom^ mice. Quantification of isolated cardiomyocytes showed ∼90%, ∼74% and ∼16% rAAV transduction rate, respectively (Fig. 4A). Echocardiogram showed that in the mid- and low-dose treated group, FS and LVID phenotypes are no longer detectable (Fig. 4B-C). The increased cardiac fibrosis and the upregulation of *Ctgf* and *Nppa* in CASAAV-treated hearts were also circumvented in the low-dose group (Fig. 4D-F), indicating this treatment allows analysis of the *Lmna* mutant cell phenotypes in a grossly normal physiological microenvironment.

**Figure 4.**
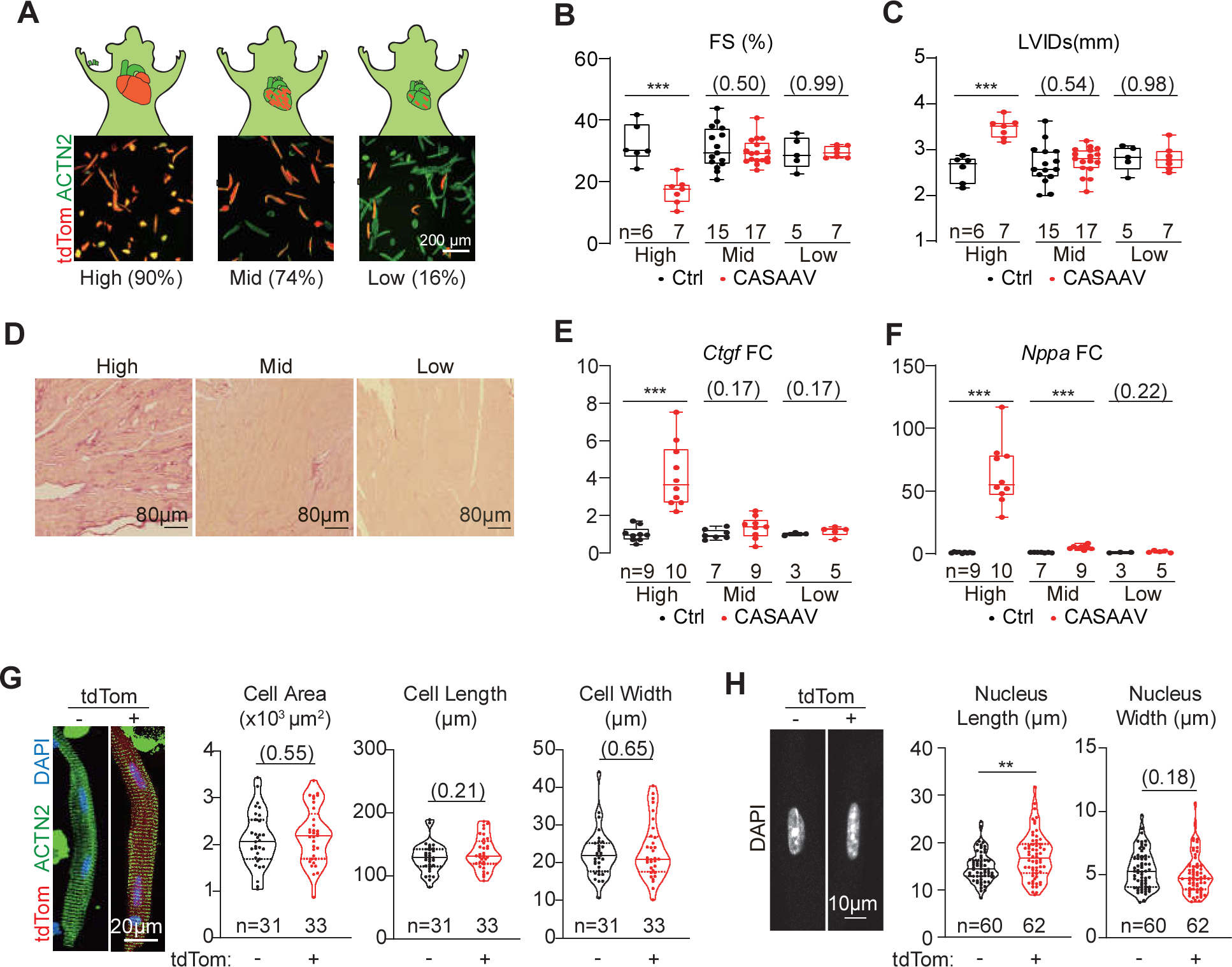
No cardiac morphology changes in *Lmna* cardiac mosaics. **(A)** Fluorescence images of cardiomyocytes that were isolated from mice treated with serial dilutions of the CASAAV vector. **(B-C)** Echocardiogram analysis of the 6-week-old mice. **(D)** Picrosirius red staining on 6-week-old heart sections. **(E-F)** RT-qPCR analysis of *Ctgf* and *Nppa*. **(G)** Immunostaining of ACTN2 on isolated cardiomyocytes and quantification of cardiomyocyte morphology. **(H)** Quantification of nuclear morphology. Two-tailed t-test: *p<0.05; **p<0.01; ***p<0.001; non-significant P values in parentheses.

We next isolated cardiomyocytes from the low-dose-treated mosaic hearts and compared the tdTom-positive versus tdTom-negative cells. We found cardiomyocyte morphology parameters were not statistically different (Fig. 4G), although the nuclei in the mutant cardiomyocytes appeared elongated (Fig. 4H). These data indicate that *Lmna* regulates cardiomyocyte morphology via a non-cell-autonomous mechanism. By contrast, *Lmna* regulates nuclear shape in a cell-autonomous manner.

### Whole-body lamin-A supplementation partially restores defects in *Lmna*^Δ/Δ^ hearts

Next we attempted to establish an AAV-based gene supplementation strategy to mitigate the cardiac defects in *Lmna*^Δ/Δ^ mice. We first constructed an AAV9 vector that delivers GFP-tagged Lamin-A protein that is expressed by the constitutive CMV promoter (AAV-CMV-LA) (Fig. 5A). A single dose of 2×10^11^ vectors was first subcutaneously injected to wildtype P1 mice to validate the transgene expression at P14 (Fig. S4A). Fluorescence imaging of heart and liver sections confirmed the nuclear periphery localization of the GFP-Lamin-A proteins (Fig. S4B). Western blot using an antibody against lamin-A/C detected the GFP-Lamin-A fusion proteins at the level that was slightly lower than the endogenous Lamin-A/C proteins (Fig. S4C). Histological analysis revealed moderate inflammatory cell infiltration into the liver of mice treated with this dose of AAV-CMV-LA for 6 months (Fig. S4D), indicating that the AAV dosage should not be further increased for safety concerns.

**Figure 5.**
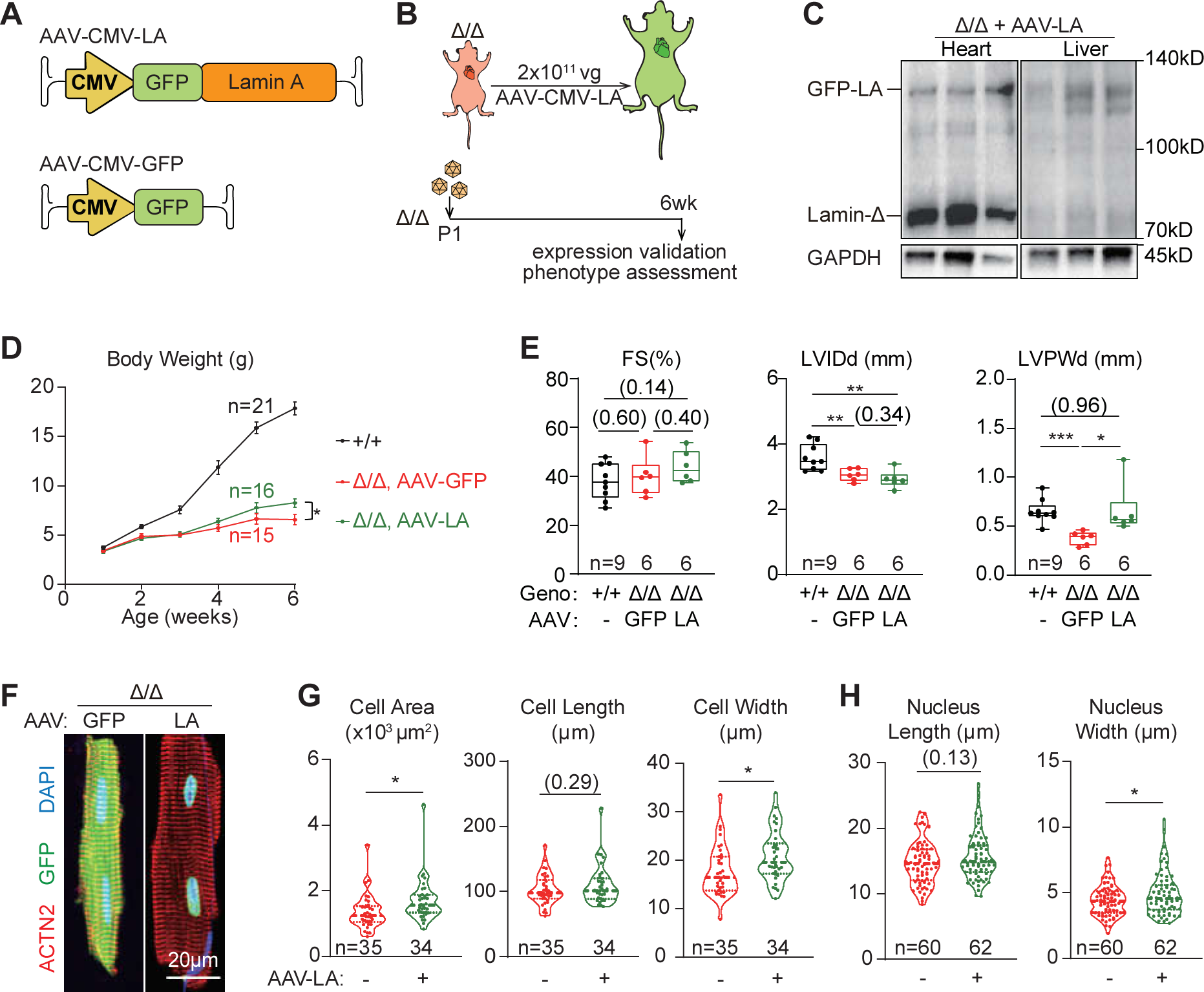
Partial rescue of cardiac morphology via whole-body lamin-A addback. **(A-B)** A diagrams showing the design of whole-body lamin-A addback experiments. **(C)** Western blot analysis of 6-week-old *Lmna*^Δ/Δ^ tissues that were treated with AAV-CMV-LA at P1. **(D)** Growth curve of animals in the whole-body lamin-A addback experiments. **(E)** Echocardiogram analysis of the 6-week-old mice. **(F-G)** Immunostaining of ACTN2 on isolated cardiomyocytes and quantification of cardiomyocyte morphology. **(H)** Quantification of nuclear morphology. Two-tailed t-test: *p<0.05; **p<0.01; ***p<0.001; non-significant P values in parentheses.

We next injected 2×10^11^ AAV-CMV-LA vectors into *Lmna*^Δ/Δ^ P1 mice to study the impact of this treatment on cardiac defects by 6 weeks after injection (Fig. 5B). Western blot validated the expression of this transgene in heart and liver (Fig. 5C). Comparing to AAV-CMV-GFP treated control animals, AAV-CMV-LA treatment moderately improved the body weight of *Lmna*^Δ/Δ^ mice (Fig. 5D). Echocardiogram revealed no changes in cardiac FS and LVID, but the ventricle wall thickness of *Lmna* ^Δ/Δ^ mice was brought back to the value comparable to wildtype controls (Fig. 5E). AAV-CMV-LA treatment improved the width and the projected area of isolated *Lmna*^Δ/Δ^ cardiomyocytes as compared to AAV-CMV-GFP treatment (Fig. 5F-G). The nuclear elongation phenotype was also alleviated by increasing the width of the cell nuclei (Fig. 5H).

We also performed a genetic mosaic analysis by comparing GFP-positive versus GFP-negative cardiomyocytes that were isolated from the same *Lmna*^Δ/Δ^ heart treated with AAV-CMV-LA (Fig. S4E). No statistic difference in cardiomyocyte morphology was detected between these two groups of cells (Fig. S4F), but GFP-positive nuclei indeed exhibited partially rescued morphology as compared to GFP-negative nuclei (Fig. S4G). These results indicate that AAV-CMV-LA improves cardiomyocyte morphology via a non-cell-autonomous mechanism, but restores nuclear morphology in a cell autonomous manner.

### Lamin-A supplementation in non-myocytes restores cardiomyocyte defects in *Lmna*^Δ/Δ^ hearts

The non-cell-autonomous rescue of cardiac defects by AAV-CMV-LA suggested that cardiomyocytes are not the correct targets for *LMNA* gene therapy. To test this idea, we first constructed an AAV-Tnnt2-LA vector in which the Tnnt2 promoter expressed Lamin-A specifically in cardiomyocytes (Fig. 6A). A single dose of 2×10^11^ vectors to wildtype P1 mice resulted in transgene expression at P14 at the level similar to endogenous lamin-A/C (Fig. 6B). When injected into *Lmna*^Δ/Δ^ animals, this dose of AAV-Tnnt2-LA added back GFP-lamin-A into ∼68% nuclei in myocardium (Fig. 6C).

**Figure 6.**
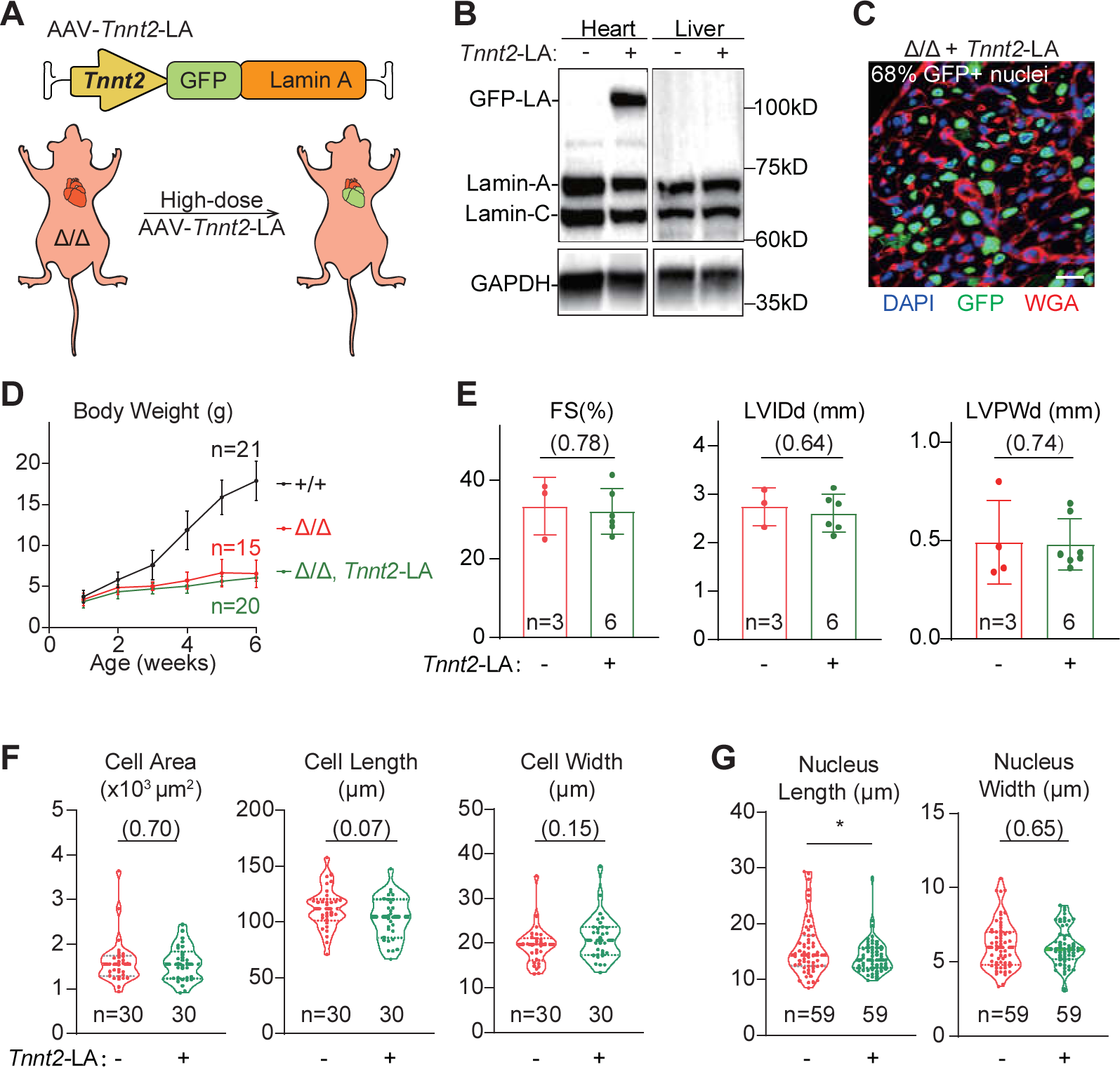
No rescue of cardiac morphology via cardiomyocyte-specific lamin-A addback. **(A)** A diagram showing the design of cardiomyocyte-specific lamin-A addback experiments. **(B)** Western blot analysis of P14 *Lmna*^Δ/Δ^ tissues that were treated with AAV-Tnnt2-LA at P1. **(C)** A fluorescence image of a P14 *Lmna*^Δ/Δ^ cardiac section that was treated with AAV-Tnnt2-LA at P1. Bar, 20μm. **(D)** Growth curve of animals in the cardiomyocyte-specific lamin-A addback experiments. **(E)** Echocardiogram analysis of the 6-week-old mice. **(F)** Quantification of cardiomyocyte morphology. **(G)** Quantification of nuclear morphology. Two-tailed t-test: *p<0.05; **p<0.01; ***p<0.001; non-significant P values in parentheses.

As compared to control *Lmna*^Δ/Δ^ animals that did not receive this AAV treatment, AAV-Tnnt2-LA-treated *Lmna*^Δ/Δ^ animals did not exhibit improved body weight (Fig. 6D), LVPW thickness (Fig. 6E) or cardiomyocyte morphology (Fig. 6F), although the nuclear elongation phenotypes were partly restored (Fig. 6G). We further administered a lower dose (1×10^10^) of the AAV-Tnnt2-LA vector into *Lmna*^Δ/Δ^ animals and created a genetic mosaic in which only 7% nuclei were GFP-LA positive (Fig. S5A-B). Comparison between the GFP-LA-positive versus GFP-LA-negative cells further confirmed the lack of cardiomyocyte rescue in *Lmna* ^Δ/Δ^ hearts upon cardiomyocyte-specific addback of lamin-A (Fig. S5C-D).

We then studied if supplementation of lamin-A into non-myocytes would be sufficient to restore cardiomyocyte defects in *Lmna*^Δ/Δ^ mice. To achieve this goal, we constructed an AAV-Loxp-CMV-Loxp-LA (AAV-Loxp-LA) vector in which the CMV promoter was flanked by Loxp sites. When this AAV was injected into mice carrying a cardiomyocyte-specific *Myh6*-Cre transgene, the CMV promoter would be deleted in cardiomyocytes, allowing GFP-lamin-A to be only expressed in non-myocyte (Fig. 7A).

**Figure 7.**
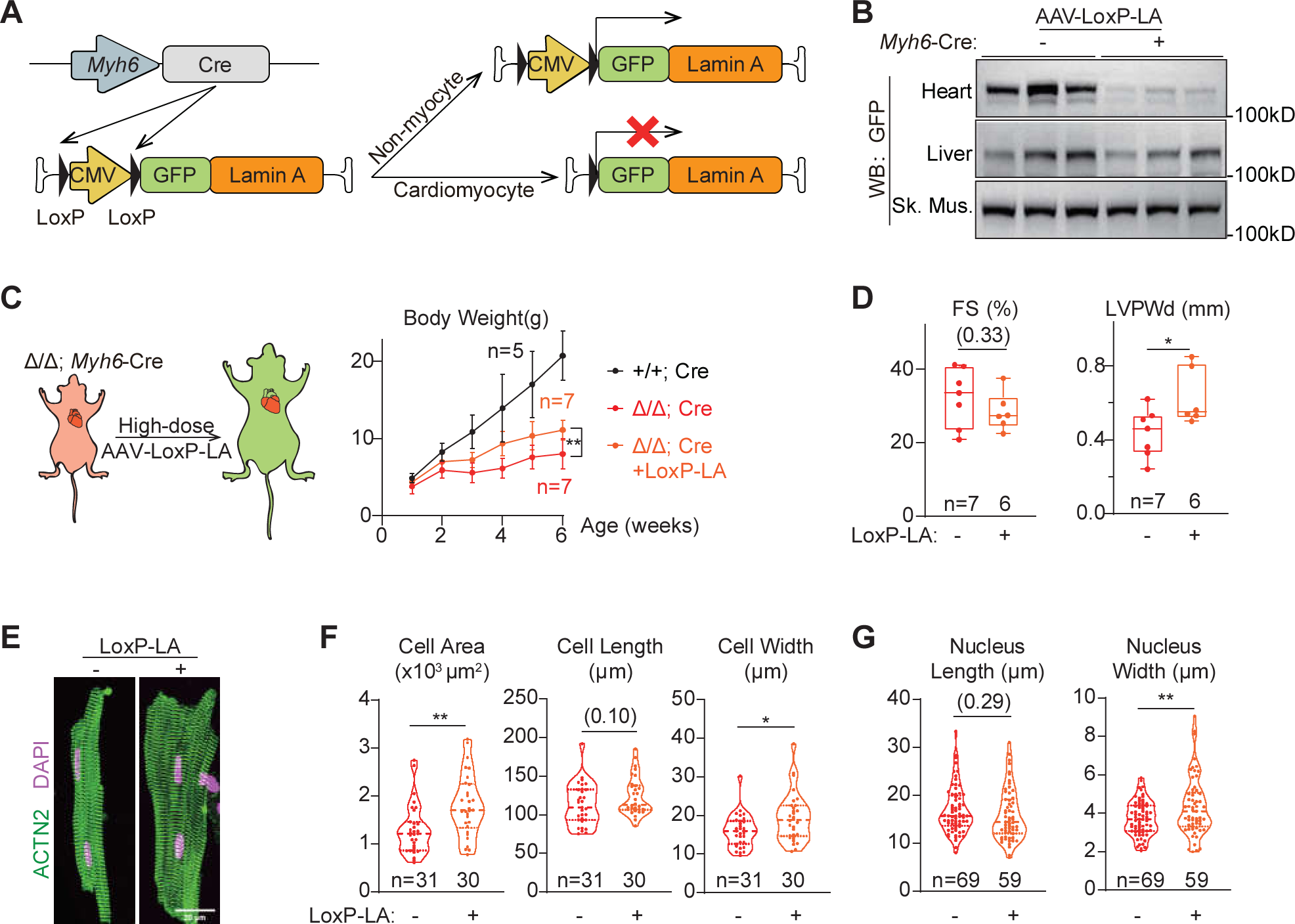
Partial rescue of cardiac morphology via cardiomyocyte-exclusive lamin-A addback. **(A)** A diagram showing the principle to achieve cardiomyocyte-exclusive lamin-A addback. **(B)** Western blot analysis of P14 tissues that were treated with AAV-Loxp-LA at P1. **(C)** Growth curve of animals in the cardiomyocyte-exclusive lamin-A addback experiments. **(D)** Echocardiogram analysis of the 6-week-old mice. **(E)** Immunostaining of ACTN2 on isolated cardiomyocytes. **(F)** Quantification of cardiomyocyte morphology. **(G)** Quantification of nuclear morphology. Two-tailed t-test: *p<0.05; **p<0.01; ***p<0.001; non-significant P values in parentheses.

We administered 2×10^11^ AAV-LoxP-LA vectors into *Lmna*^Δ/Δ^;Tg(Myh6-Cre) mice at P1 and assessed transgene expression at P14. As compared to *Lmna*^Δ/Δ^ mice that do not carry the Myh6-Cre allele, immunofluorescence (Fig. S6A-B) and western blot analysis demonstrated reduced GFP-LA signal in the heart of *Lmna*^Δ/Δ^;Tg(Myh6-Cre) mice while the transgene expression in liver and skeletal muscles were not affected (Fig. 7B and Fig. S6C). Interestingly, AAV-LoxP-LA treatment significantly improved body weight growth and LVPW thickness in *Lmna*^Δ/Δ^;Tg(Myh6-Cre) mice (Fig. 7C-D). Cardiomyocyte and nuclei morphology was also partially restored (Fig. 7E-G) in the *Lmna*^Δ/Δ^; Tg(Myh6-Cre) mice treated with AAV-LoxP-LA. Together, these data show that non-myocytes are the primary cell types in which Lamin-A supplementation exert a therapeutic effect on cardiomyocyte defects.

## Discussion

In this study, we introduced *Lmna* truncating mutations in three different forms of mouse models to understand the mechanisms by which *Lmna* mutations cause cardiomyocyte defects. In mice carrying the mutation throughout the body, atrophic cardiac phenotypes were observed. By contrast, cardiac specific mutagenesis at the same genetic loci resulted in cardiac hypertrophy. These seemingly contradictory phenotypes could be reconciled via a cardiac genetic mosaic analysis, which unambiguously indicated that *Lmna* did not perform a cell-autonomous function in cardiomyocyte morphology regulation. Instead, the atrophic phenotype in germline mutants is secondary to global developmental defects, while the hypertrophic phenotype in cardiomyocyte specific mutants is secondary to cardiac systolic dysfunction. Thus, this study highlights genetic mosaic analysis as a powerful and necessary approach to precisely define molecular mechanisms in cardiac pathogenesis, circumventing secondary effects that could lead to misunderstanding of gene functions in diseases.

Previous studies harnessed either germline or cardiac specific mutagenesis approaches to establish animal models of *LMNA* DCMs, but these two approaches were never directly compared to determine how they represent different aspects of this disease. In this study, we showed that cardiomyocyte-specific *Lmna* truncating mutations triggered severe DCM phenotypes while germline mutants could maintain normal cardiac systolic functions, as evaluated by fraction shortening measurement. These distinct cardiac phenotypes indicate that DCM pathogenesis is not only a consequence of myocardial weakness, but also influenced by the imbalance between cardiac maturation versus the growth of the body. In the germline mutants, although the heart is atrophic, cardiac dysfunction is largely compensated by the global deficiency in animal development. These results indicate that it is necessary to systemically compare multiple models and investigate the whole body in addition to the heart in order to fully understand DCM pathogenesis.

The contribution of non-myocytes, potentially the cells in non-heart tissues, to *LMNA* DCM pathogenesis is also supported by our attempt to establish an AAV gene supplementation therapy. We decided to use germline mutants, but not cardiac specific mutants, to test this idea because most *LMNA* DCM patients carry the mutation throughout the body. Several types of non-myocytes have indeed been recently reported to contribute to *LMNA* DCM pathogenesis^21, 22^. In sharp contrast to the established notion that cardiomyocyte is the primary cell type in which *LMNA* mutation causes DCM, supplementation of lamin-A specifically in cardiomyocytes does not rescue the cardiac growth phenotypes in germline mutants. By contrast, cardiomyocyte-excluded lamin-A addback is sufficient to restore cardiac growth similar to the results of whole-body lamin-A supplementation. These data indicate that non-myocytes are essential targets in gene therapy for *LMNA*-associated cardiac defects.

It is important to note that lamin-A supplementation by rAAV rescues nuclear morphology phenotypes, regardless whether lamin-A addback is achieved throughout the body or specifically in cardiomyocytes. These results agree to the well-established notion that lamin-A cell-autonomously determines nuclear mechanics, which lays the basis for nuclear shape regulation. Together, the distinct mechanisms by which *Lmna* regulates cardiomyocyte morphology versus nuclear morphology indicate a requirement of simultaneously adding back *LMNA* to both non-myocytes and myocytes in gene therapy.

A major unsolved problem in this study is to identify the key cell type that contributes to AAV-based mitigation of cardiac phenotypes in the germline mutants. One group of candidate cells include the major non-myocytes in the heart, such as endothelial cells^22^ and fibroblasts^21^. However, the contribution of these cells to AAV therapy is probably low due to the limited transduction of AAV9 into the non-myocytes in the heart. Another group of cell types that could potentially contribute to cardiac therapy include cells in non-heart organs. For example, in the AAV-LoxP-LA-based gene supplementation experiments, lamin-A was efficiently added back to the liver and skeletal muscles. Further investigation of these organs in *LMNA* gene therapy could potentially uncover inter-organ communications as new pathogenic mechanisms underlying *LMNA* DCM pathogenesis, and establish novel therapeutic strategies for these diseases.

Our study also demonstrated the precise control of AAV dosage and *LMNA* transgene expression level as another challenge in AAV-*LMNA* therapy. We carefully titrated AAV-CMV-LA but we were not able to identify a high AAV dose that enables transgene expression to be at the same level of the endogenous gene in the heart. By contrast, in our safety assessment, AAV transduction and transgene expression causes detectable adverse effects in the liver, confirming the hepatic injury as a major problem in gene therapy involving high AAV dosage^38, 39^.

Overall, this study indicates that a successful gene supplementation therapy requires careful dissection of cell-autonomous versus non-cell-autonomous causes of the disease. This investigation is necessary to determine the cell targets for gene therapy, which is not necessarily the same cell type that primarily presents disease phenotypes.

## Materials and Methods

(See supplementary information for more information about materials and methods.)

### Mice

All procedures involving experimental animals were performed in accordance with protocols approved by the Institutional Animal Care and Use Committee of Peking University, China (approval number LA2021332), and conformed to the Guide for the Care and Use of Laboratory Animals (8th edition. The National Academies Press, 2011) by the Association for Assessment and Accreditation of Laboratory Animal Care.

*Lmna*^Δ/Δ^ mice were generated while we were generating another *Lmna* mutant as previously described^31^ at the Institute of Laboratory Animal Science, Chinese Academy of Medical Sciences, via CRISPR/Cas9-based zygotic mutagenesis. The sgRNA sequences used in generating this allele and the genotyping primer sequences were shown in Figure S3. The Rosa^Cas9-Tom^ mice were purchased from GemPharmatech (Strain No. T002249). The *Myh6-Cre* mice were purchased from Cyagen Biosciences (Strain No. C001041).

All mice were kept in a temperature-controlled room (21 ± 1°C) with a 12-hour light/dark cycle and had free access to water and normal chow. After genotyping at postnatal day 0, neonatal *Lmna*^Δ^*^/^*^Δ^ mice under inhalation anesthesia by isoflurane were administrated with rAAVs or vehicle subcutaneously. When necessary, adult mice were euthanized by cervical dislocation.

### Adult cardiomyocyte isolation

Cardiomyocytes were isolated by retrograde perfusion^27^. In brief, heparin-treated mice were anesthetized with 3% isoflurane. Hearts were extracted and cannulated onto a Langendorff perfusion apparatus. 37 °C perfusion buffer was first pumped into the heart to flush out blood and equilibrate the heart. Collagenase II (Worthington, LS004177) was next perfused into the heart for 8 min at 37 °C to dissociate cardiomyocytes. The apex was cut from the digested heart, gently dissociated into single cardiomyocytes in 10% FBS/perfusion buffer and filtered through a 100 µm cell strainer to remove undigested tissues. To prepare for immunostaining, cardiomyocytes were cultured in DMEM+10%FBS medium for 30min on a laminin-coated cover glass before fixation with 4% PFA and permeabilization with 4%BSA+0.1% TritonX-100/PBS.

### RNA sequencing analysis

The libraries were constructed using NEBNext Ultra RNA Library Prep Kit for Illumina (E7530L, NEB). Sequencing was performed on an Illumina NovaSeq 6000 platform with 2D×D150 bp pair-end reads at Novogene, China. RNA-seq reads were aligned to mm10 by STAR^40^ and reads counts were calculated by FeatureCounts^41^. DESeq2^42^ was used to perform statistical analysis of differential gene expression. An adjusted P value of 0.05 was used as the cutoff to identify differentially regulated genes. GO term analysis was performed using GSEA analysis with ranked gene lists^43^.

### Amplicon sequencing

Genomic DNA was extracted from tissues using TIANamp Genomic DNA Kit (DP304, Tiangen, China). The sgRNA-targeted loci of *Lmna* gene were amplified using Taq PCR MasterMix (KT211, Tiangen, China) and purified by TIANgel Purification Kit (DP219, TIANGEN). See Table S3 for primer sequences, which include all sequences necessary for sequencing library construction. Sequencing was performed on an Illumina NovaSeq 6000 platform with 150 bp single-end reads at Novogene, China. The sequencing results were processed by CRISPResso2^44^. The output bam files were transformed into sam files by samtools view v1.6^45, 46^ and exploited to calculate the frameshifting indel rates of the target regions using a home-made python script.

### AAV design and production

The human lamin-A coding sequence was acquired at Addgene (#124268) and subcloned into the AAV-Tnnt2-GFP-v2^30^ (addgene#165036, control vector in this study) to build the AAV-Tnnt2-GFP-LA plasmid. Subsequently, the Tnnt2 promoter was replaced by the CMV promoter or the LoxP-CMV-LoxP cassette to generate AAV-CMV-GFP-LA and AAV-LoxP-CMV-LoxP-GFP-LA plasmids, respectively. These new plasmids will be available at Addgene soon after publication. CASAAV plasmids were produced as previously described^27^. Among the two sgRNAs used to target *Lmna*, one sgRNA is identical to the one used to generate the *Lmna*^Δ^ allele^31^.

AAV production was performed in house or at PackGene Biotech. We produced AAV9 as previously described^27, 36^. In brief, 140 µg AAV-ITR, 140 µg AAV9-Rep/Cap, and 320 µg pHelper (pAd-deltaF6, Penn Vector Core) plasmids were produced by maxiprep (DP117, Tiangen, China), and triple transfected into HEK293T cells in ten 15-cm plates. 60-72 h after transfection, cells were resuspended in lysis buffer (20 mM Tris pH=8, 150 mM NaCl, 1 mM MgCl_2_, 50 µg/ml Benzonase) and lysed by two freeze-thaw cycles. AAV in culture medium was precipitated by PEG8000 (VWR, 97061-100), resuspended in lysis buffer and pooled with cell lysates. AAV particles were next purified in an Optiprep density gradient (D1556, Sigma) by ultracentrifugation (Beckman Optima XPN-100) with a type 70Ti rotor. The AAV were next concentrated in PBS with 0.001% pluronic F68 (Invitrogen, 24040032) using a 100 kD filter tube (Fisher Scientific, UFC910024). AAV titer was quantified by real-time quantitative PCR using a fragment of the GFP coding sequence to make a standard curve. See Table S3 for primer information.

### Statistical Analysis

Statistical analysis and plotting were performed using GraphPad Prism. Statistical tests are indicated in each figure legend. Numbers in parentheses in figures indicate non-significant P-values. Bar plots show mean ± standard deviation. In box plots, horizontal lines indicate the median and 25th and 75th quantiles; whiskers extend to the extreme values; dots represent raw data points. Statistical analyses were performed using Student’s unpaired t test. P values less than 0.05 were considered to indicate statistically significant differences.

### Data and materials availability

Next generation sequencing data have been deposited at National Genomic Data Center (https://ngdc.cncb.ac.cn/gsa/) under the accession code CRA011594 (amplicon-sequencing), CRA011590 (6-week *Lmna*^Δ/Δ^ and *Lmna*^+/+^ RNA-seq) and CRA011582 (2-week *Lmna*^+/+^ RNA-seq). All AAV plasmids will be available at Addgene soon after publication. All full-length raw western blots are available in Figure S7. Other data supporting the findings of this study are available upon reasonable request.

## Abbreviations

AAV: adeno-associated virus
DCM: dilated cardiomyopathy
LOF: loss-of-function

## Acknowledgements

We thank the Institute of Laboratory Animal Science at Chinese Academy of Medical Sciences for the production of *Lmna*^Δ/Δ^ mice. We thank PackGene Biotech for service in AAV production.

## Author Contribution

Y.G. conceived the research and wrote the paper. Y.G., X.H. and S.Z. supervised the execution of this project. W.T.P. and B.J. provided supports in AAV-Tnnt2-LA and CASAAV vector construction and scientific advice. E.D. and M.Z. provided technical assistance. Y.S. and Z.C. performed RNA-Seq analysis. D.Z. and C.W. conducted amplicon-seq data analysis. Y.Z. assisted in figure preparation and graphic illustration. C.G., J.L., and L.Y. assisted in animal husbandry and characterization. Y.S. conducted experiments, collected data and organized the results.

## Ethics Statement

All animal strains and procedures were approved by the Institutional Animal Care and Use Committee of Peking University.

