## Supplementary Information for "Non-cell autonomous cardiomyocyte regulation complicates gene supplementation therapy for *LMNA* cardiomyopathy"

### 1. Supplementary Figure

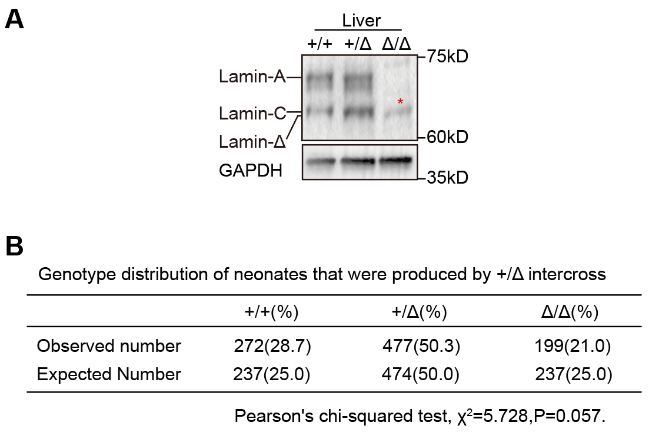

#### Figure S1. Characterization of *Lmna*^Δ/Δ^ mice.

**(A)** Western blot analysis of liver. **(B)** Chi-square analysis of genotype distribution at birth.

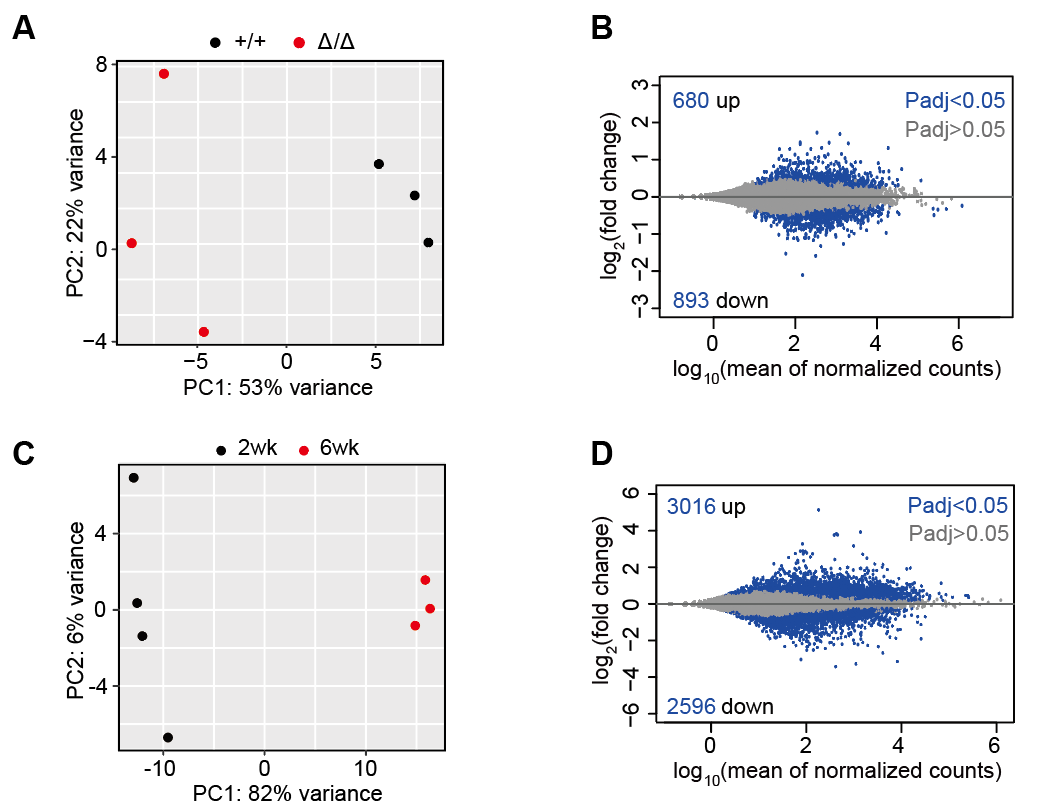

#### Figure S2. Characterization of RNA-Seq data of cardiac tissues.

**(A-B)** PCA and MA plots of 6-week *Lmna*^Δ/Δ^ versus *Lmna*^+/+^ ventricles. **(C-D)** PCA and MA plots of 6-week versus 2-week *Lmna*^+/+^ ventricles.

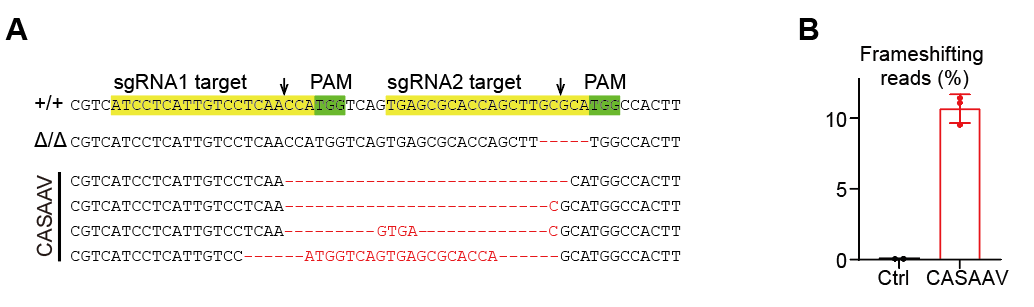

#### Figure S3. Amplicon-seq analysis of CASAAV treatment.

**(A)** Representative reads in amplicon-seq data. Red dashes indicate deleted bases. **(B)** Quantification of frameshifting mutations in amplicon-seq data.

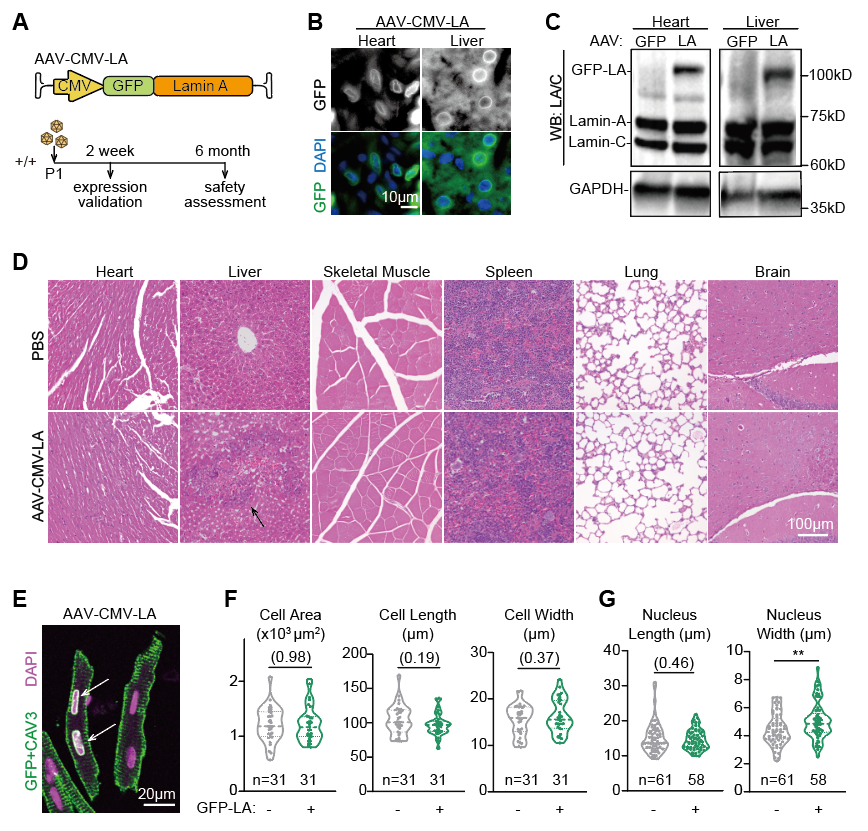

#### Figure S4. Characterization of AAV-CMV-LA vectors.

**(A)** A diagram of the AAV assessment experiment in wildtype mice. **(B)** Fluorescence images of AAV-CMV-LA treated tissues at P14. **(C)** Western blot analysis of Lamin-A/C in AAV-CMV-LA treated tissues at P14. **(D)** H&E staining of tissues that were collected from mice treated with AAV-CMV-LA for 6 months. Arrow indicates accumulation of inflammatory cells in the liver. **(E)** Immunofluorescence of isolated cardiomyocytes. Arrows indicate the GFP-LA-positive nuclei in a GFP-positive cell to the left. A GFP-negative cells is at the right. **(F-G)** Quantification of cellular **(F)** and nuclear **(G)** morphology. Two-tailed t-test: **p<0.01; non-significant P values in parentheses.

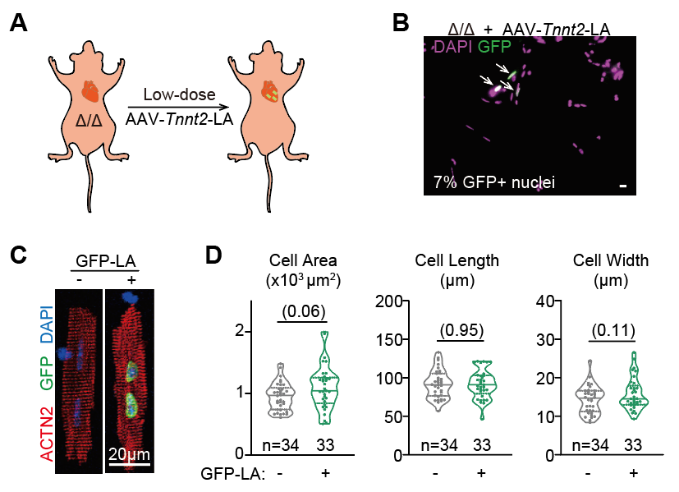

#### Figure S5. Mosaic lamin-A addback experiment.

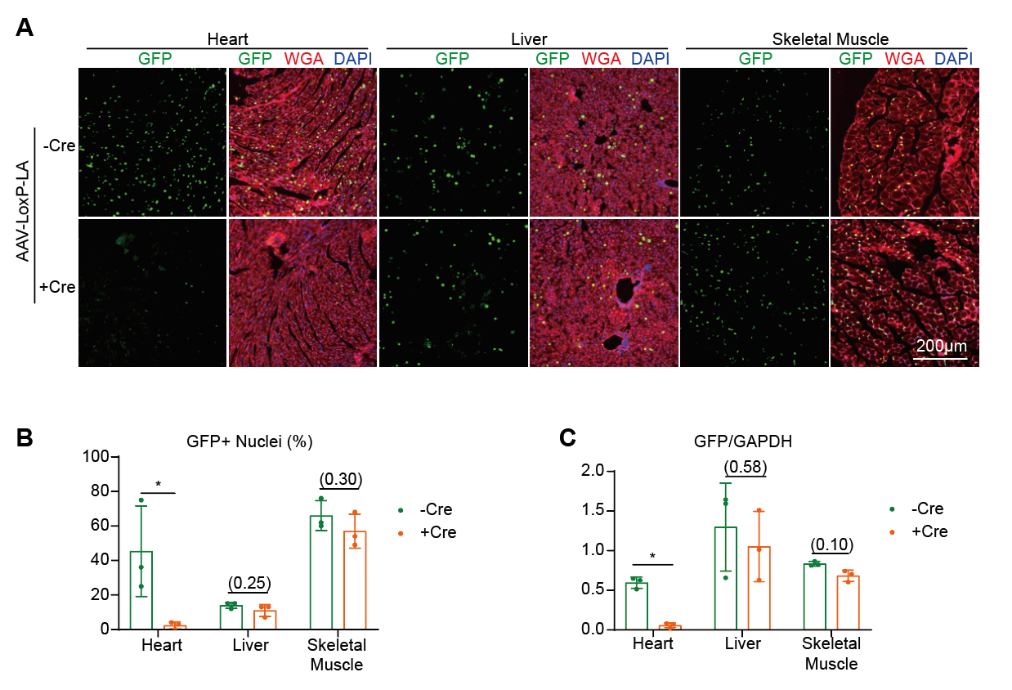
**(A)** A diagram showing cardiomyocyte-specific lamin-A addback in a genetic mosaic fashion. **(B)** Fluorescence imaging of nuclei in isolated cardiomyocytes. Arrows indicate GFP-LA-positive nuclei. Bar, 20μm. **(C)** ACTN2 immunostaining of GFP-positive and GFP-negative cardiomyocytes. **(D)** Cell morphology quantification. Two-tailed t-test: Non-significant P values in parentheses.

#### Figure S6. Validation of cardiomyocyte-excluded lamin-A addback.

**(A)** Fluorescence imaging of tissue sections that were treated with AAV-Loxp-LA vectors. **(B)** Quantification of GFP-positive nuclei in tissue sections. **(C)** Quantification of GFP in western blot analysis. Two-tailed t-test: *p<0.05; non-significant P values in parentheses.

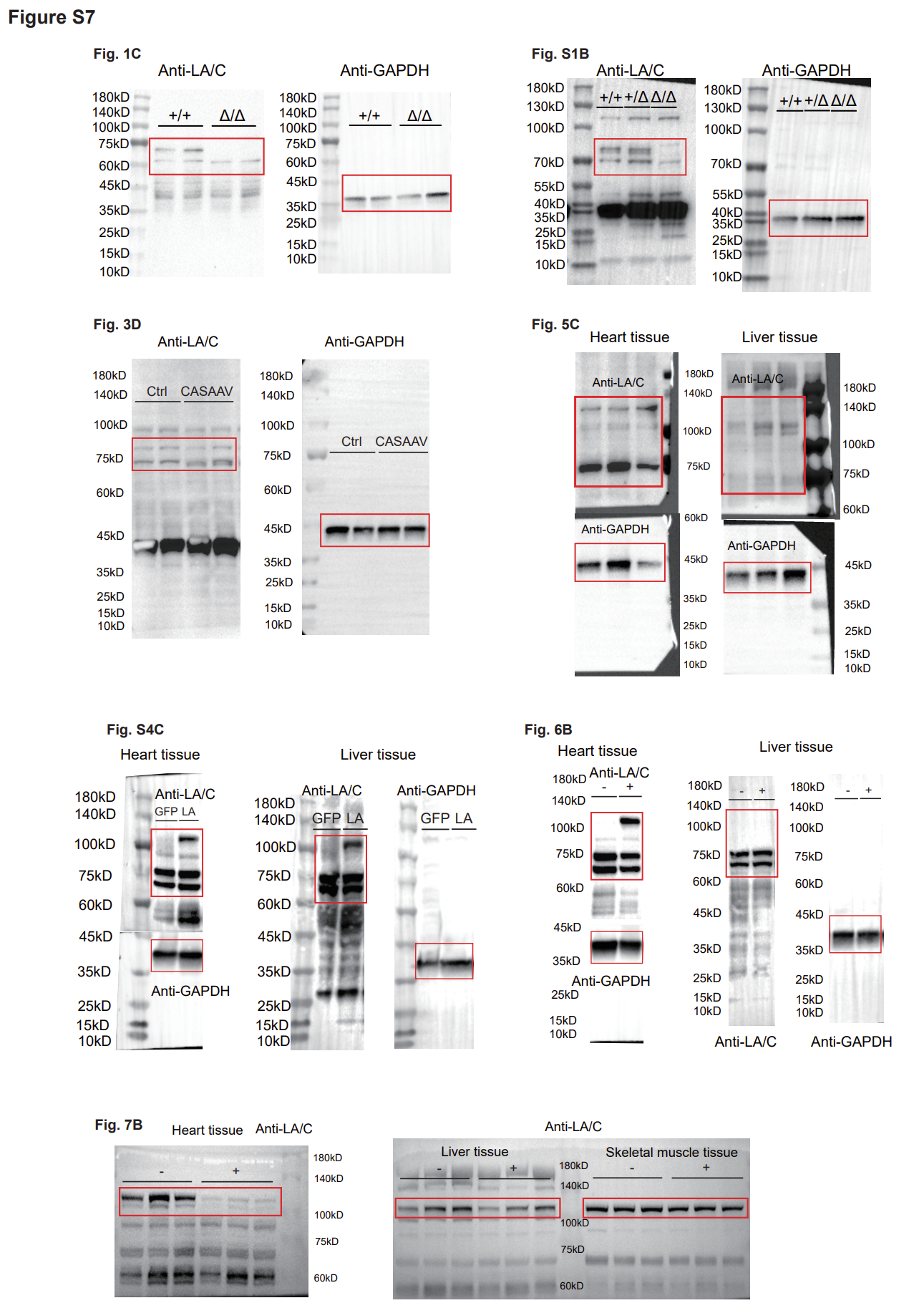

#### Figure S7. Full-length original western blot images.

### 2. Supplementary Table

#### Table S1. RNA-Seq differential expression analysis of 6-week *Lmna*^Δ/Δ^ versus *Lmna*^+/+^ ventricles (file separately attached)

#### Table S2. RNA-Seq differential expression analysis of 6-week versus 2-week *Lmna*^+/+^ ventricles (file separately attached)

#### Table S3. DNA primers in this study

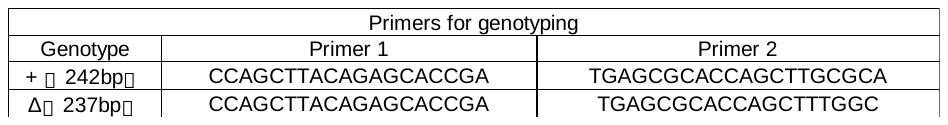

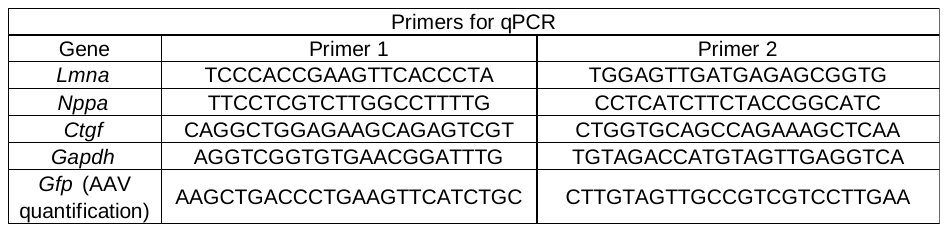

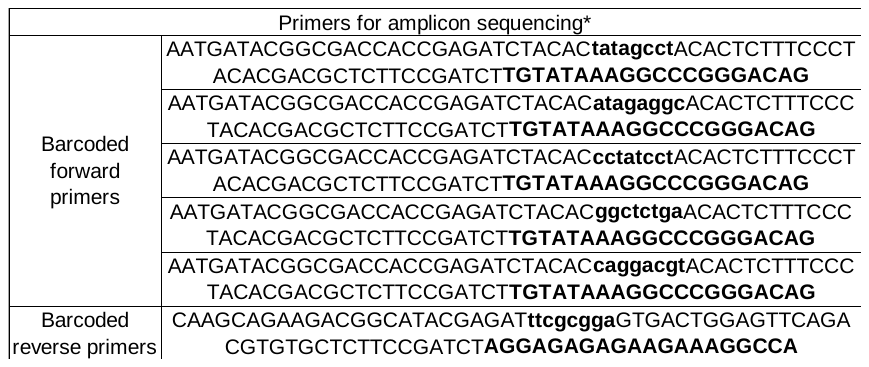

*barcodes and genome-matching sequences in bold.

#### Table S4. Antibodies and dyes in this study

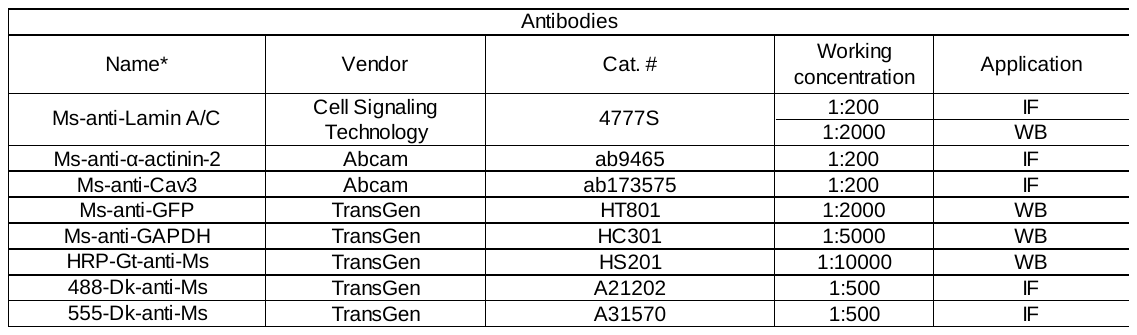

*Ms: mouse, Gt:goat, Dk:donkey

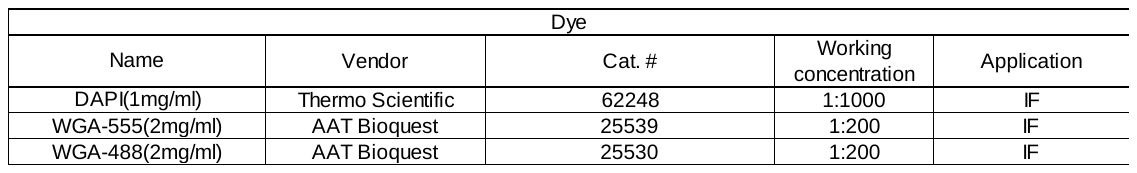

### 3. Supplementary Materials & Methods

#### Echocardiography

Echocardiography was measured on mice that were anesthetized initially by 3% isoflurane and maintained asleep at 1~1.5% isoflurane (R510-22, RWD, China). Echocardiography was performed with a VINNO6n machine (VINNO Corporation, Suzhou, China). The standard left ventricular short-axis cardiac videos were acquired using a 23MHz transducer under M mode. FS, LVPW and LVID values were measured by averaging results from five consecutive heart beats. The researcher who performed echocardiography was blinded to the genotypes of the animals.

#### Tissue sectioning

Paraffin sectioning was performed at Servicebio, China. Heart tissues were first fixed in 4% paraformaldehyde (PFA) at 4°C overnight and then in 20% sucrose solution overnight. Next, the samples were dehydrated through a serial gradient of ethanol and N-butanol. The samples were waxed in liquid paraffin for 2 h and embedded to make paraffin blocks using a tissue embedder (EG1150, Leica, Germany). Four-micron sections were cut by paraffin slicer (RM2245, Leica, Germany) and adhered to the microscope slides after floating on a water bath (HI1210, Leica, Germany).

For frozen sectioning, the hearts were fixed with 4% PFA and were dehydrated in sucrose solutions (15% followed by 30%) overnight. Then the tissue samples were embedded in OCT (Sakura) and frozen in -80°C. Heart sections (7 μm) were cut on a cryostat microtome (CM1950, Leica, Germany).

#### Histological staining

Histological staining was performed at Servicebio using paraffin sections. In brief, for H&E staining, the paraffin sections were dewaxed in xylene, hydrated by passing decreasing concentrations of alcohol baths and then stained in 0.5% hematoxylin (G1003, Servicebio) for 3-5min. Then the sections were washed in running water and differentiated by dipping in 1% acid ethanol for a few seconds. Next the sections were rinsed in water, and stained in 0.05% eosin Y (G1003, Servicebio) for 10min. Then the sections were dehydrated in increasing concentrations of ethanol, cleared in xylene and mounted for imaging.

For picrosirius red staining, the paraffin sections were dewaxed and dehydrated and then incubated with 0.2% picrosirius red solution dissolved in saturated aqueous picric acid (1.2% picric acid in water) (G1018, Servicebio). Next the sections were rinsed in water, dehydrated in increasing concentrations of ethanol, cleared in xylene and mounted for imaging.

#### Western blotting

The tissues was washed once on ice phosphate buffered saline (PBS). Then the tissues were lysed in RIPA Buffer (25mM Tris PH7.0~8.0, 150mM NaCl, 0.1% SDS, 0.5% sodium deoxycholate, 1% Triton X-100) containing protease inhibitors (Solarbio, China). Tissue lysates were centrifuged for 15 min at 1,2000 rpm at 4°C. The supernatants were collected and the protein concentration was quantified using BCA Protein assay kit (TransGen Biotech, Beijing, China).

The lysates were diluted into identical protein concentrations, added into 4×SDS sample buffer and heated at 70℃ for 10 min. Then 30 μg proteins of each sample were subjected to SDS-PAGE in a 4-12% gradient gel (TransGen Biotech, China) using a constant voltage at 120 V for 1 h 30 min. The proteins were transferred to a PVDF membrane and blocked for 1 h in 5% milk/TBST at room temperature. The primary antibodies were incubated with the membranes overnight at 4°C. After the membranes were washed with TBST, the HRP-conjugated secondary antibodies were used in staining for 1 h at RT. After washing, the ECL Western blotting substrate (PE0010, Solarbio, Beijing, China) was used to detect chemiluminescence signals with a Bio-Rad imager (Bio-Rad, USA). See Table S4 for antibody information.

#### RNA extraction and quantitative real-time PCR (RT-qPCR)

Total RNA was extracted from heart apex using the TransZol Up Plus RNA Kit (ER501-01, TransGen, Beijing, China). TransScript II One-Step gDNA Removal and cDNA Synthesis SuperMix (AH311-03, TransGen, Beijing, China) kits were used for genomic DNA removal and reverse transcription. Real-time PCR analysis was performed using Perfect Start Green qPCR Super Mix (+DyeII) (AQ602-24, TransGen, Beijing, China) and the AriaMx Real-Time PCR System (Agilent Technologies). See Table S3 for primer information.

#### Immunofluorescence

The samples were incubated with primary antibodies overnight at 4°C. Then the samples were washed with PBS for 3 times, 5 min each. Fluorescent secondary antibodies and dyes were incubated with the samples for 1 h at room temperature. Then the slides were mounted with ProLong™ Diamond antifade mountant (P36961, Invitrogen, Thermo Fisher Scientific, USA). See Table S4 for antibody and dye information.

Epifluorescence images were acquired using an all-in-one fluorescence microscope (BZ-X800, Keyence, Japan). The confocal images were observed by an SP8 (Leica, Germany) confocal microscope equipped with an APO 63X/1.4 oil objective. ImageJ was used for quantitative measurement. Cell morphology, Z-line distance and nuclear morphology parameters were manually measured.
